# Evidence for a regulatory role of the evolutionarily conserved sequence complementarity between tRNAs

**DOI:** 10.64898/2026.09.02.748885

**Authors:** Amit Kumar Sahu, Arpan Amlanjyoti Dash, Umesh Varshney

## Abstract

The tRNAs are non-coding RNAs (ncRNAs) known for their classical role in decoding mRNAs. Analysis of tRNA sequences from *Escherichia coli* revealed pairwise similarities for different combinations of tRNAs, as well as complementarities between some tRNA pairs. To explore the physiological significance of the complementarities between the tRNAs, we investigated the pair of tRNAs encoded by *lysT* (encoding an abundant tRNA, tRNA^LysT^), and *argU* (encoding a rare tRNA, tRNA^ArgU^). We show that tRNA^LysT^ and tRNA^ArgU^ anneal to form a heterodimer *in vitro*. The heterodimerisation is prevented by the presence of DNA oligomers complementary to the interacting sequences, in a dose dependent manner. Consistent with the notion of sequestration of tRNA^ArgU^ by tRNA^LysT^, while the overexpression of tRNA^ArgU^ did not impact the culture growth, that of tRNA^LysT^ did. The tRNA^LysT^ mediated inhibition of the culture growth was enhanced at a lower temperature. The AGA minigene (decoded by tRNA^ArgU^) mediated toxicity and hybrid phage (λ_imm_-P22) growth on Δ*ssrA* (tmRNA) strains was also consistent with the sequestration of tRNA^ArgU^ by tRNA^LysT^. Also, we observed tRNA-derived fragments (tRFs) from tRNA^LysT^ and tRNA^ArgU^, which too might facilitate tRNA^ArgU^ sequestration. Taken together, these observations support a novel regulatory role of the evolutionary conserved complementarities between tRNAs.

**Importance:** Besides carrying amino acids to ribosomes, tRNAs play important non-canonical roles in cells. Also, the tRNA-derived fragments (tRFs) contribute to the non-canonical roles of tRNAs. Analysis of tRNA sequences from *Escherichia coli* reveals evolutionarily conserved pairwise complementarity between tRNAs. We investigated the importance of the complementarity between tRNA^LysT^ (an abundant tRNA) and tRNA^ArgU^ (a rare tRNA). The tRNA^LysT^ and tRNA^ArgU^ heterodimerise *in vitro*. And, consistent with the notion of sequestration of tRNA^ArgU^ by tRNA^LysT^, overexpression of tRNA^LysT^, inhibits the culture growth, enhances the AGA minigene toxicity, and growth of the hybrid phage (λ_imm_-P22) on Δ*ssrA* (tmRNA) strains, supporting a novel regulatory role of the evolutionary conserved complementarities between tRNAs. The tRNA-derived fragments (tRFs) may also contribute to this role.

## Introduction

Transfer RNAs (tRNAs) are the most abundant RNA molecules after the rRNAs in the cells. The primary function of the tRNAs is to decode the codons in mRNAs into polypeptide chains during translation by bringing cognate amino acids to the ribosome. The tRNAs show the highly conserved cloverleaf secondary structure and the L-shaped tertiary structure, stabilized by extensive intramolecular base-pairing and post-transcriptional modifications (Cramer et al., 1969; Holley et al., 1965; Väre et al., 2017). The folding of tRNA is essential for its recognition by aminoacyl-tRNA synthetases, translation factors, and the ribosome.

Although tRNAs are conventionally known as adaptor molecules during the translational process, growing evidence supports their participation in other cellular processes such as amino acid synthesis, tetrapyrrole synthesis, modifications of peptides lipids, viral genome replication, viral or mobile genetic element insertion in the genome, and other tRNA dependent regulatory functions (Katz et al., 2016; Kirchner & Ignatova, 2014). Also, the tRNA-derived fragments (tRFs) have roles in regulating gene expression, stress responses, and cellular adaptation (Muthukumar et al., 2024; Shen et al., 2018). These studies suggest that tRNAs engage in molecular interactions that extend beyond their canonical role in protein synthesis.

The stability of the secondary/tertiary structures of a tRNA, formed through intramolecular base pairing, depend on multiple factors such as the chaperone proteins, nucleoside modifications, ionic strength, and the kinetics of transcription and folding (Bhaskaran et al., 2014; Keffer-Wilkes et al., 2016a, 2020). This suggests that tRNAs may obtain alternative conformations and intermolecular interactions under certain environmental conditions. An interesting but less explored possibility is that different tRNAs containing complementary sequences may be involved in direct tRNA-tRNA interactions and assist formation of homodimeric or heterodimeric tRNA duplexes. In fact, some of the earlier studies have shown the formation of tRNA dimers (Loehr & Keller, 1968; Yang et al., 1972), suggesting that these interaction may have physiological significance (Tosar et al., 2018). In more recent times, tRNA-tRNA interactions have been reported using advanced analysis technique (Liu et al., 2017). However, little is known about the frequency, nature, and biological relevance of these interactions.

To investigate the possibility of intermolecular interactions among bacterial tRNAs, we carried out pairwise sequence comparisons of all tRNAs expressed in *E. coli* using the RNAup server. This study revealed multiple pairs of tRNAs with significant sequence similarities due to their shared evolutionary origins and conserved structural necessities. Interestingly, we also discovered several pairs of tRNAs exhibiting extensive regions of sequence complementarity. These observations indicated that specific tRNAs may have an inherent ability of participating in the direct intermolecular base-pairing interactions. We hypothesized that tRNAs might transiently unfold to allow formation of intermolecular complexes with the complementary tRNAs under suitable cellular conditions. These interactions may affect tRNA availability, stability, aminoacylation, or the translation process, offering an unidentified mechanism of post-transcriptional regulation.

To explore the functional significance of such potential dimerization in bacterial genetic system, we focused on a pair of tRNAs comprising a rare tRNA encoded by *argU* (tRNA^ArgU^), and an abundant tRNA encoded by *lysT* (tRNA^LysT^). Both of these tRNAs share a significant stretch of sequence complementary, making them attractive candidates for intermolecular interaction. Our findings provide evidence of previously unrecognized phenomenon of tRNA– tRNA interactions and suggest that heteromerizations of tRNAs could represent a novel layer of translational regulation in cells.

## Results

### Sequence complementarities between tRNAs

Comparisons of tRNA sequences from *E. coli* that are available from the genomic tRNA database (GtRNAdb) revealed stretches of intermolecular complementarities (**Table S1**). The extent of the complementarity depended on the allowed mismatches or the gaps. The tRNA^ArgU^ and tRNA^LysT^ pair (**Fig. 1A**) from *E. coli* MG1655 showed significant complementarity across their TΨC and the acceptor arms (**Fig. 1B**). As tRNA^ArgU^ is a rare tRNA, we used the tRNA^ArgU^ and tRNA^LysT^ pair as a test case to investigate the importance of the sequence complementarities between the two tRNAs. We then performed RNAup analysis to evaluate the accessibility-corrected interaction potential between tRNA^LysT^ and each of the remaining annotated tRNAs. Interaction energies varied between -8.64 and -2.55 kcal/mol (**Table S2**) with the tRNA^ArgU^ and tRNA^LysT^ pair showing one of the highest favorable value (ΔG = −8.64 kcal/mol).

**Fig. 1:**
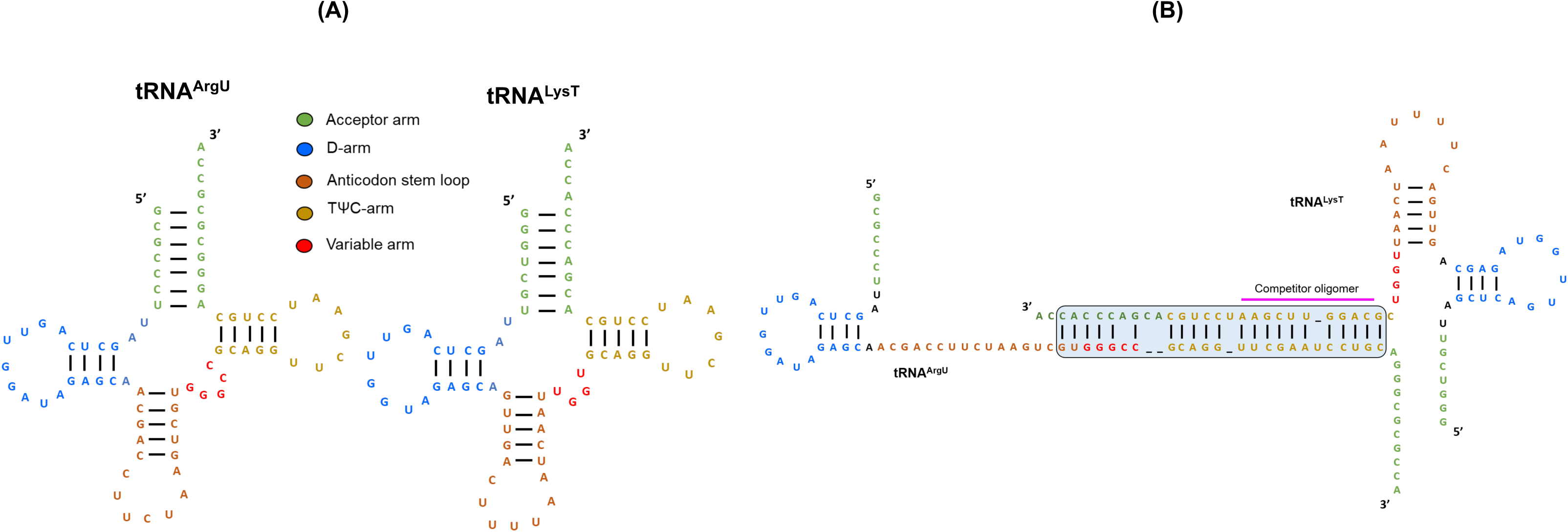
**(A)** Cloverleaf structure of tRNA^LysT^ and tRNA^ArgU^, depicting different arms of the tRNA with different colors. Green-acceptor arm, Blue-D arm, Light brown-anticodon arm, Red-variable loop, and mustard yellow-TΨC arm. **(B)** Schematic representation of tRNA^LysT^ and tRNA^ArgU^ extended interaction due to the presence of complementary regions between TΨC and acceptor arm of tRNA^LysT^ and tRNA^ArgU^ with few mismatches. The region where the competitor oligomer (represented as pink in colour) binds is shown in the diagram.

### tRNA^LysT^ and tRNA^ArgU^ interact *in vitro*

To explore the relevance of the complementarity between tRNA^ArgU^ and tRNA^LysT^, we performed tRNA interaction/annealing experiments using their *in vitro* transcribed RNA and mixing them in equal concentrations (**Fig. 2A**), and analysing them on the native PAGE. We observed a band of higher molecular weight only in the reactions where both tRNA^ArgU^ and tRNA^LysT^ were present (**Fig. 2B**, lanes 4 and 5, compare with lanes 1 and 2). Given that the same band appeared when either tRNA^LysT^ (**Fig. 2B**, lane 4) or tRNA^ArgU^ (**Fig. 2B**, lane 5) was used as radiolabelled probe, it indicated that the two tRNAs interacted/heterodimerised. To check the specificity of this interaction, we used tRNA^ValU^, which did not show any significant complementarity with either the tRNA^LysT^ or tRNA^ArgU^, as control. Under the same conditions, we did not observe any bands corresponding to interaction/heterodimerisation of either the tRNA^LysT^ or tRNA^ArgU^ with the tRNA^ValU^ (**Fig. 2B**, lanes 6 and 8, compare with lanes 1-3).

**Fig. 2:**
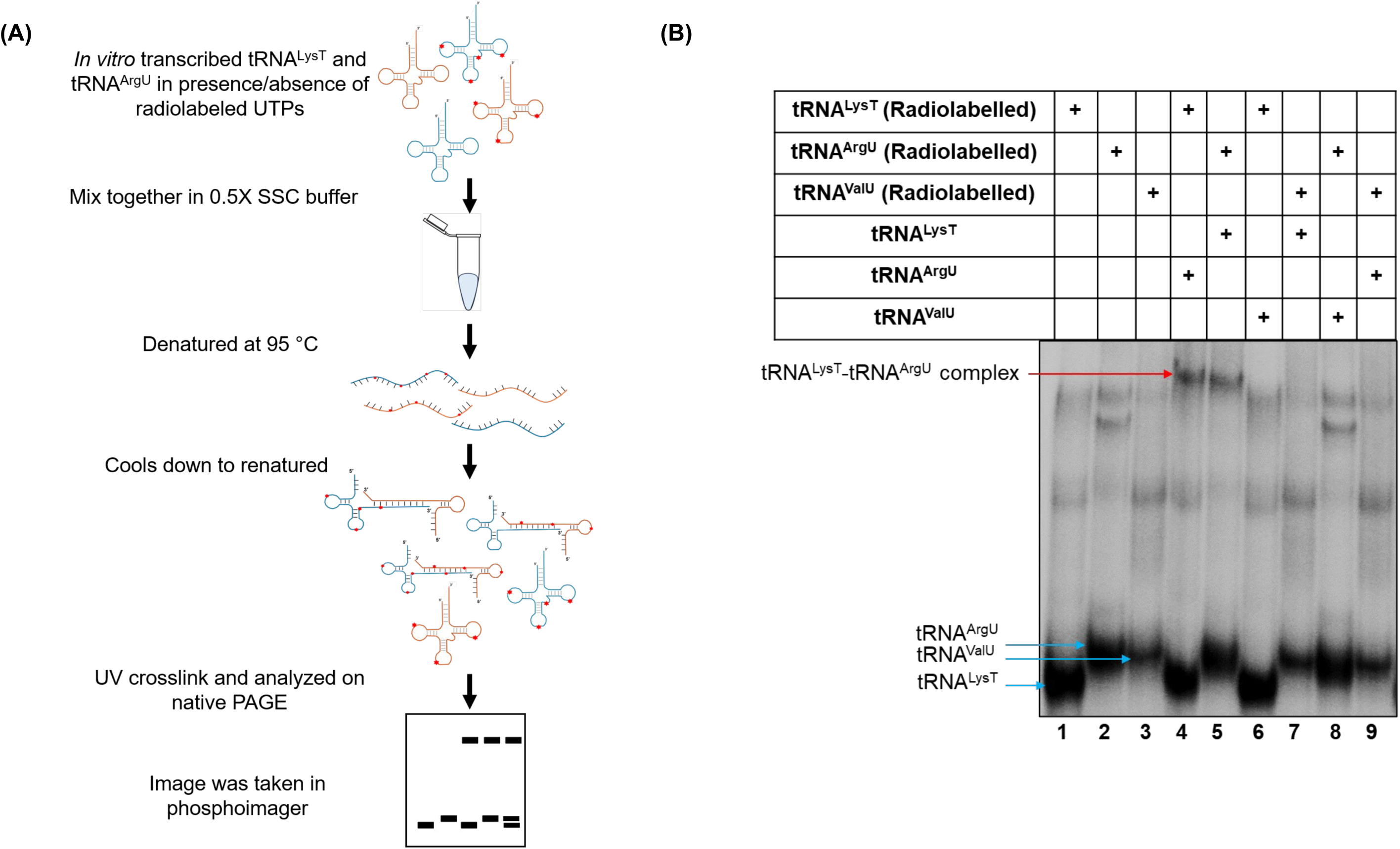
*In vitro* reaction shows tRNA^LysT^ and tRNA^ArgU^ interact with each other and form dimer. **(A)** Schematic representation of protocol followed for *in vitro* tRNA binding assay. The reaction was carried out with radiolabelled *in vitro* transcribed tRNA in 0.5X SSC buffer. **(B)** Radiogram showing the tRNA^LysT^-tRNA^ArgU^ complex. The tRNA heterodimer band is marked by red arrow. The tRNA monomers are marked by blue arrow.

### tRNA^LysT^ and tRNA^ArgU^ form heterodimer at specific complementary sites

*In silico* analysis showed the presence of complementary sequences between tRNA^ArgU^ and tRNA^LysT^. To verify if the binding/heterodimerisation between tRNA^ArgU^ and tRNA^LysT^ occurred because of a specific interaction between the predicted complementary regions, we repeated the experiment in the presence of a competitor anti-tRNA^ArgU^ or anti-tRNA^LysT^. DNA oligomers that bind to the complementary regions between the two tRNAs (**Fig. 1B**), were used in the assay and the reactions were analyzed by native PAGE followed by northern blotting (**Fig. 3A**). We observed that the extent of the tRNA heterodimer complex formation gradually declined with the increasing concentration of the competitor oligomer (**Figs. 3B** and **3C; Fig. S1**). As the tRNA concentrations were kept the same in all the reactions, these observations suggest that the interaction between the two tRNAs is specific and mediated by the presence of the predicted complementary sequences.

**Fig. 3:**
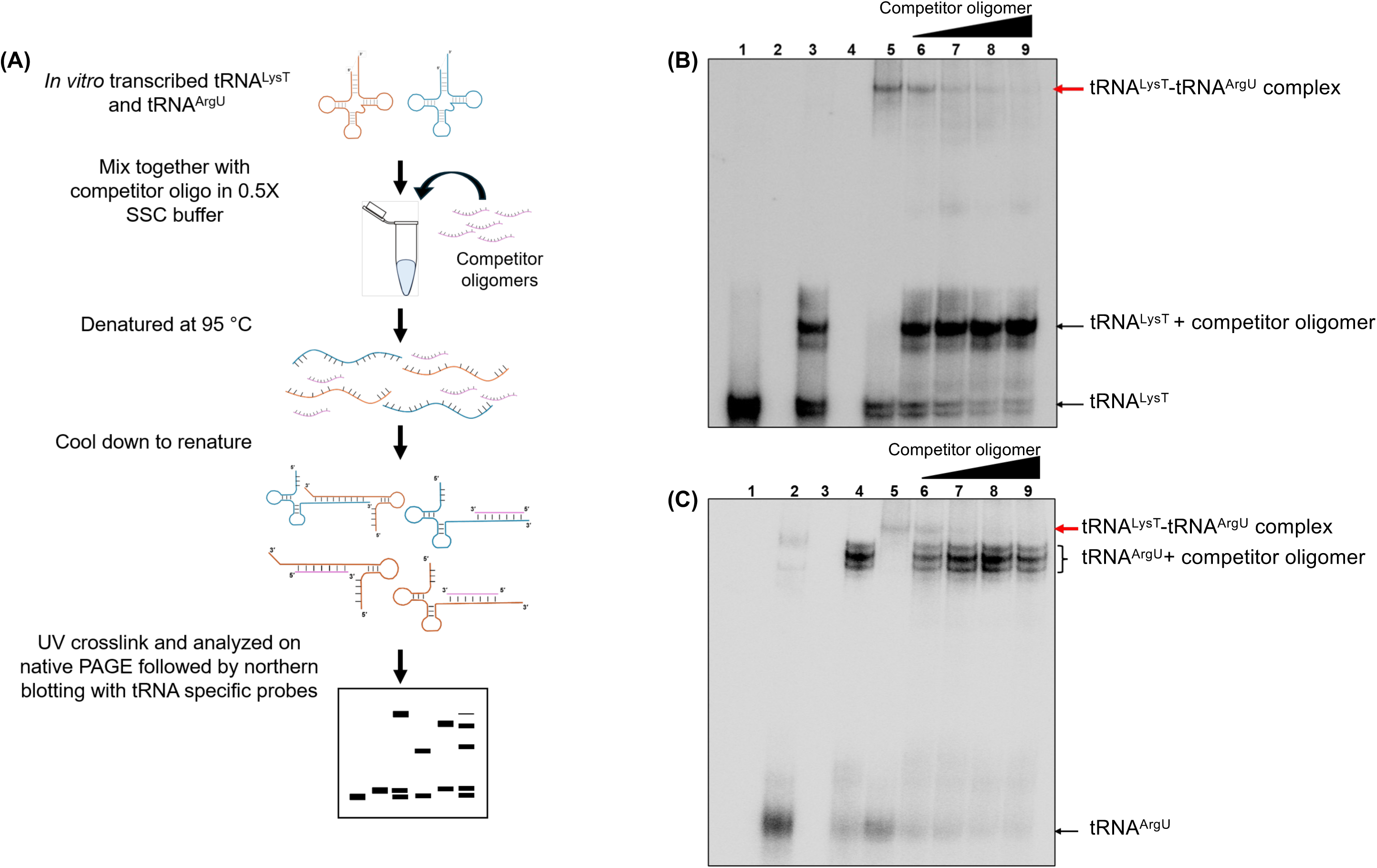
**(A)** Schematic representation of protocol followed for *in vitro* tRNA binding in the presence of competitor DNA oligomer. Northern blots depicting bands for tRNA^LysT^ **(B)** and tRNA^ArgU^ **(C). Lanes:** 1, tRNA^LysT^; 2, tRNA^ArgU^; 3, tRNA^LysT^ + competitor oligomer; 4, tRNA^ArgU^ + competitor oligomer; 5, tRNA^LysT^ + tRNA^ArgU^; 6-9, tRNA^LysT^ + tRNA^ArgU^ + competitor oligomer. Both panels show decrease in tRNA dimer formation between tRNA^LysT^ and tRNA^ArgU^ in the presence of competitor oligomer. Red arrows show the tRNA^LysT^-tRNA^ArgU^ dimer formation.

### Temperature-dependent growth variation on expression of plasmid based constructs of tRNA^LysT^ and tRNA^ArgU^

To investigate the effect of the plasmid based expression of the two tRNAs *in vivo*, we cloned the tRNA genes with their native promoters in pBAD vector to generate pBAD*lysT* and pBAD*argU* constructs (**Fig. 4A**). The pBAD, pBAD*lysT* and pBAD*argU* plasmids were then introduced into *E. coli* BW25113. The strains (vector control, *lysT* and *argU*, respectively) were checked for growth in liquid and solid media at different temperatures, without adding any arabinose (**Figs. 4B** and **4C**). We observed unremarkable growth differences between tRNA^ArgU^ and tRNA^LysT^ strains at 37 °C with their growths being similar to the vector control. However, at the lower temperature (28 °C), while no growth defect was observed for the *argU* strain, the *lysT* strain grew slower than the vector control. Similar observations were made in other media conditions (**Fig. S2**). These observations support tRNA^LysT^ toxicity on growth, especially at the lower temperature, which would favor tRNA^ArgU^ and tRNA^LysT^ interaction.

**Fig. 4:**
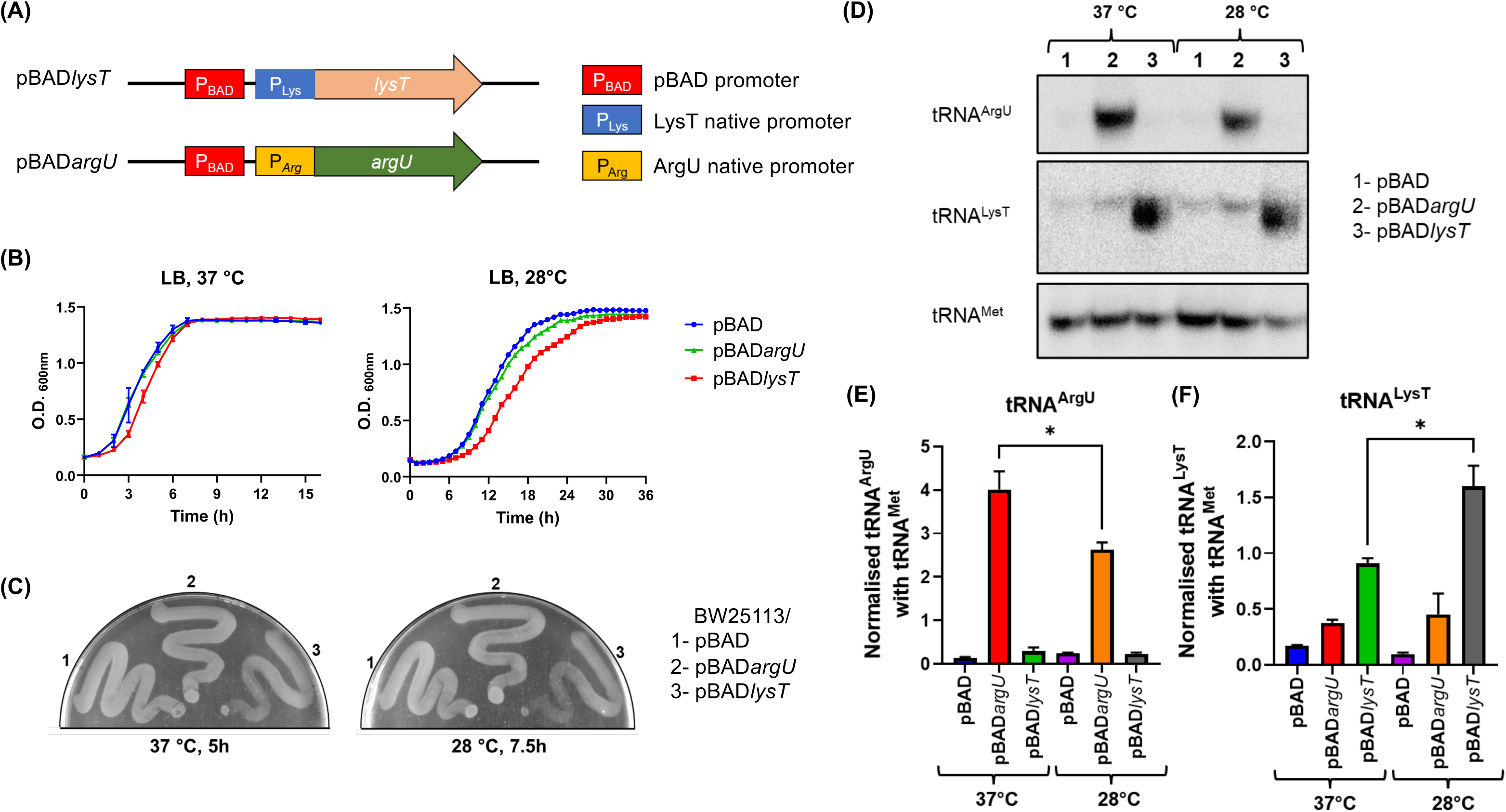
**(A)** Schematic representation of *lysT* and *argU* gene cloning in pBAD vector. Growth curve analysis **(B);** and plate assays **(C),** of strains overexpressing tRNA^LysT^ or tRNA^ArgU^ at 37 and 28 °C. The tRNA^LysT^ overexpression shows slower growth at the low temperature. **(D)** tRNA^LysT^ and tRNA^ArgU^ levels analysed by northern blotting following their separation on native PAGE. **(E-F)** Graphs showing the quantification of the tRNA levels from the northern blot. Both tRNA^LysT^ and tRNA^ArgU^ were normalised with Met elongator tRNA.

### Differences in the levels of tRNA^LysT^ and tRNA^ArgU^

For the growth decrease in the *lysT* strain, we hypothesized that an increase in the level of tRNA^LysT^ may sequester tNRA^ArgU^. Thus, a further decrease in the levels of an already a rare tRNA^ArgU^ by tRNA^LysT^ might be a cause of toxicity/slow growth in the strain. To check this, we performed northern analysis for the levels of tRNAs in the cell. We found that the relative level of tRNA^ArgU^ was higher at 37 °C than at 28 °C. Interestingly, the relative level of tRNA^LysT^ was found to be increased at lower temperature (**Figs. 4D-F**). Together with the possibility of better stability of the heteromers of the two tRNAs, these data are consistent with the notion of tRNA^ArgU^ sequestration by tRNA^LysT^.

### Impact of expression of plasmid constructs of tRNA^LysT^ and tRNA^ArgU^ on λ_imm_-P22 hybrid phage growth

The λ_imm_-P22 hybrid phage contains short translatable sequences (‘minigenes’) comprising AGA(Arg) codons in its immunity region. As the AGA codon is decoded by tRNA^ArgU^, a rare tRNA, these minigenes result in stalling of ribosomes that are rescued by trans-translation using tmRNA (transfer-messenger RNA) encoded by *ssrA* gene (Singh et al., 2008). The tmRNA contains a tRNA-like domain and a short mRNA region whose first codon (also known as resume codon) consists of Ala codon (GCA). Not unexpectedly, the λ_imm_-P22 hybrid phage fails to replicate in *E. coli sip* (*ssrA*) mutant (Retallack et al., 1994). Substitutions of the tmRNA resume codon from Ala to any rare codons reduce the trans-translation efficiency of tmRNA (Kapoor et al., 2011). We made use of this observation as an assay system to further investigate the *in vivo* impact of tRNA^ArgU^ or tRNA^LysT^ overexpression on the cellular physiology by determining the plaque forming capacity of the λ_imm_-P22 hybrid phage on the *ssrA* mutant strains. The *E. coli* BW25113 Δ*ssrA*::*kan* strains were complemented with either wild type *ssrA* (*ssrA*^WT^) or the *ssrA* where the resume codon was changed from GCA to AGA (*ssrA*^AGA^). In the phage plating experiment, we observed a good number of plaques in the *ssrA*^WT^ but very few plaques in the *ssrA*^AGA^ strains. However, when the *ssrA*^AGA^ strain was supported with excess tRNA^ArgU^, the plaque formation was rescued to the level seen on the *ssrA*^WT^ strain (**Fig. 5A**). When we introduced pBAD*lysT* in the the *ssrA*^AGA^ strain, the newly generated strain grew extremely poorly. As a control, plaque forming units were observed in tRNA^ArgU^ (*ssrA*^WT^/pBAD*argU*) or tRNA^LysT^ (*ssrA*^WT^/pBAD*lysT*) strains in the *ssrA*^WT^ background. However, the plaque sizes on *ssrA*^WT^/pBAD*lysT* strain were smaller compared to those on the *ssrA*^WT^/pBAD*argU* (**Fig. 5B**) and the same is supported by tRNA^LysT^ and tRNA^ArgU^ levels (**Fig. S3**). While the strain with the higher levels of tRNA^ArgU^ forms normal size plaques, the strain with the higher levels of tRNA^LysT^ inhibits both the plaque formation and the plaque size (**Figs. 5B** and **S3B, S3C**). Overall, these observations are also consistent with the hypothesis that both the tRNA^ArgU^ and tRNA^LysT^ interact with each other (which leads to the sequestration of tRNA^ArgU^) restricting the growth of the *ssrA*^AGA^/pBAD*lysT* strain.

**Fig. 5:**
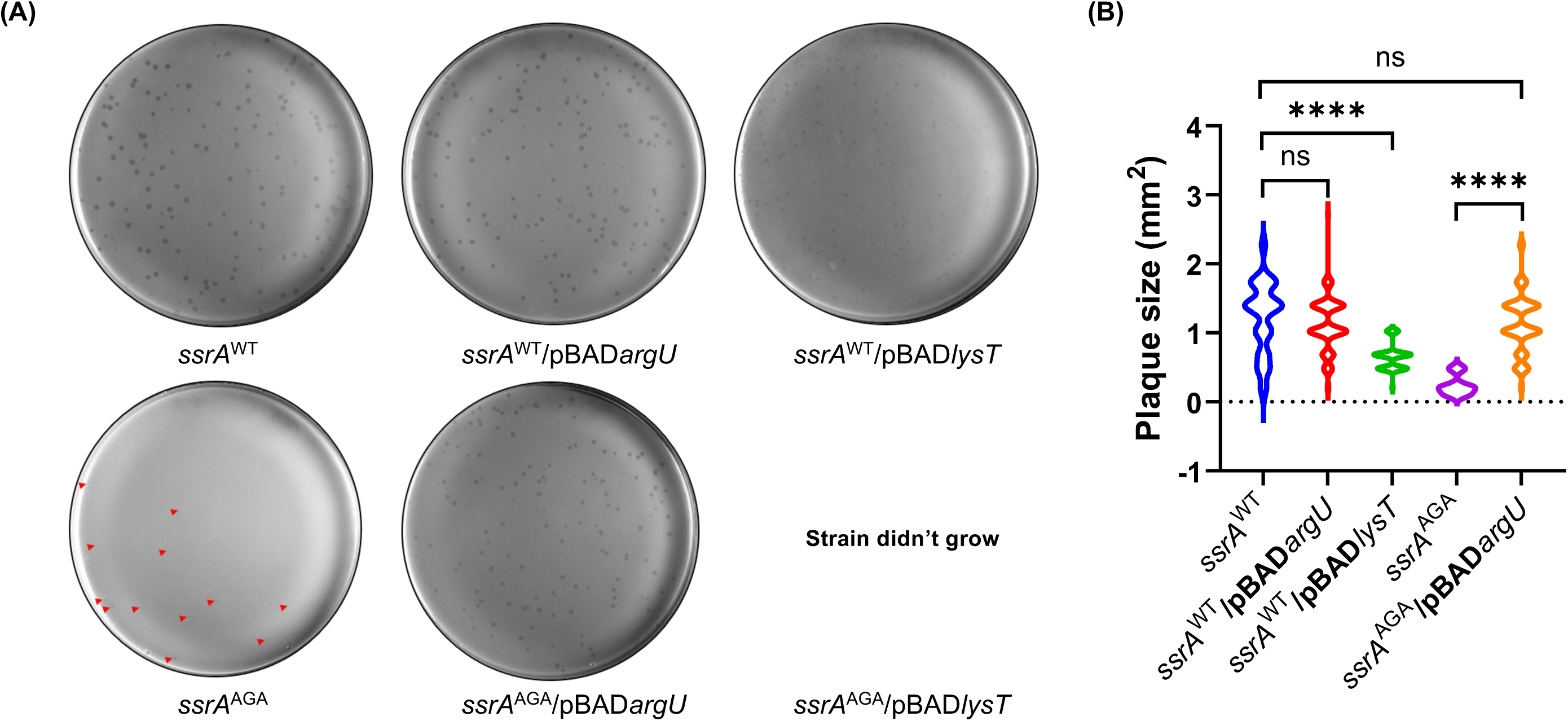
**(A)** Phage plating assay showing impact of tRNA^LysT^ and tRNA^ArgU^ overexpression on phage growth in *ssrA*^WT^ and *ssrA*^AGA^. The tRNA^LysT^ and tRNA^ArgU^ overexpression is from their native promoters. **(B)** Violin plot showing the differences in the size of plaques formed on *ssrA*^WT^ and *ssrA*^AGA^ mutant strains overexpressed with tRNA^LysT^ and tRNA^ArgU^.

### Impact of the plasmid constructs of tRNA^LysT^ and tRNA^ArgU^ on AGA minigene toxicity

To further check for the tRNA^LysT^ toxicity, we used the AGA minigene assay. The AGA codon is read by tRNA^ArgU^ and the AGA minigene has been shown to cause toxicity to *E. coli* (Cruz-Vera et al., 2003). In our earlier studies, we showed that the AGA minigene, with ORF of ATG AGA AGA TGA, sequestered tRNA^ArgU^ and caused toxicity in *E. coli* (Singh et al., 2008). We used IPTG inducible AGA minigene, and the plasmid based constructs of the tRNAs under their native promoter, cloned in the same reporter plasmids (**Fig. 6A)**. In this assay too, while tRNA^ArgU^ expression rescued the AGA minigene toxicity, the tRNA^LysT^ expression enhanced it (**Figs. 6B-E; S4**). Taken together, these observations once again support the notion that excess of tRNA^LysT^ affects tRNA^ArgU^ function.

**Fig. 6:**
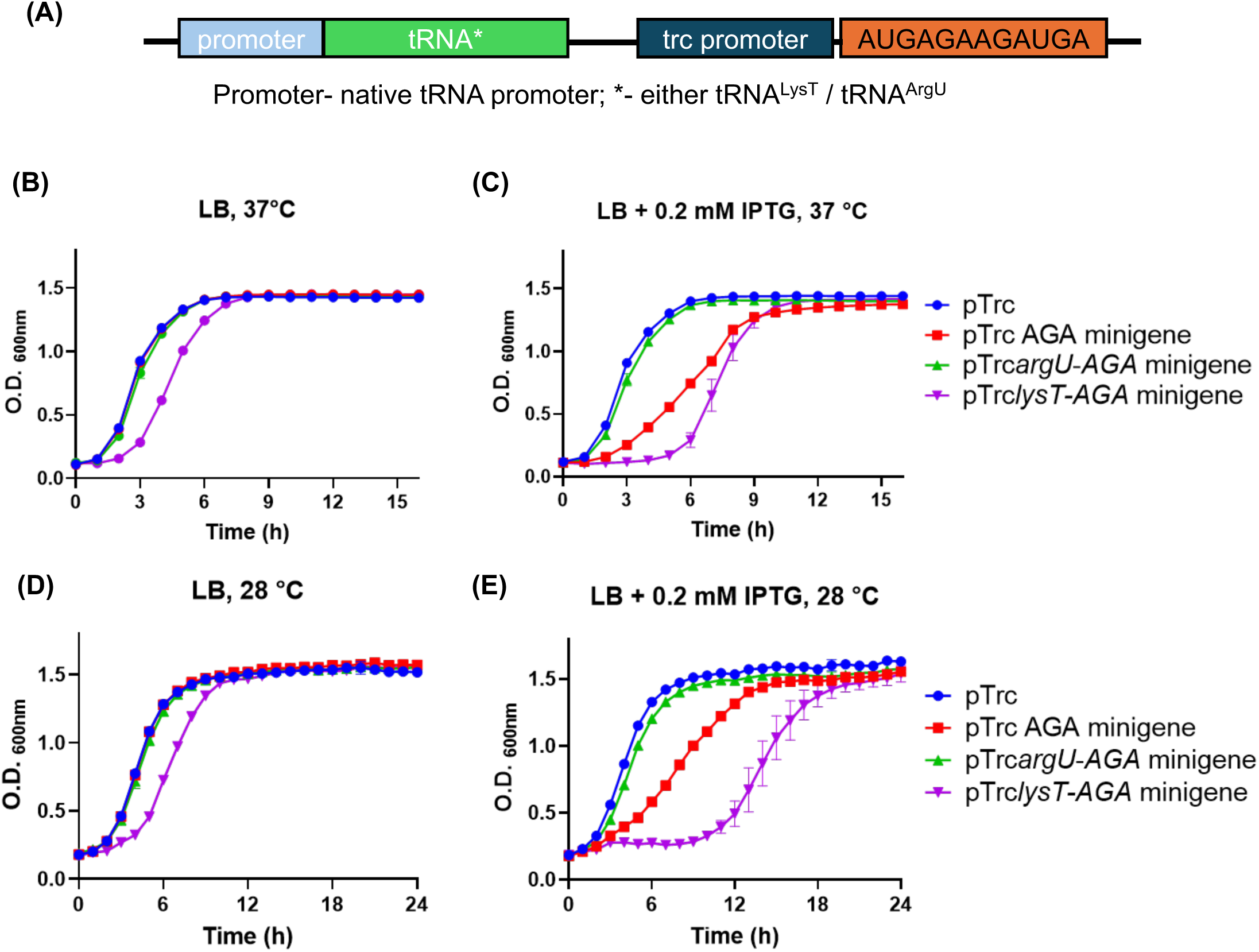
The AGA minigene reporter provides evidence of tRNA^LysT^-tRNA^ArgU^ interaction *in vivo*. **(A)** Schematic representation of tRNA gene and *AGA* minigene arrangement on the reporter plasmid. While *AGA* minigene expression is under the IPTG inducible promoter, plasmid based expression of tRNAs is under their native promoters. **(B-E)** Growth curves showing increased toxicity of AGA minigene upon tRNA^LysT^ overexpression, suggesting tRNA^ArgU^ sequestration by tRNA^LysT^.

### Both the tRNA^LysT^ and tRNA^ArgU^ form tRNA-derived fragments (tRFs)

The sequences that are complementary between the tRNA^ArgU^ and tRNA^LysT^ are already involved in the secondary structure formation of the cloverleaf. However, as also shown in **Figs. 5** and **6**, abundance of tRNA^LysT^ may still sequester a fraction of tRNA^ArgU^ into a tRNA^ArgU^-tRNA^LysT^ complexes. The tRNA-derived fragments (tRFs) have been reported even in bacteria. It is possible that the tRFs may also contribute to making tRNA^ArgU^ non-functional. To check for this possibility, we investigated for the presence of tRNA^ArgU^ or tRNA^LysT^ fragments by northern blotting using tRNA specific probes (**Figs. 7A** and **B**). The tRFs were confirmed by small RNA sequencing analysis which shows the formation of the tRNA fragments (**Fig. 7C**; **Fig. S5**). As shown in **Fig. S5**, we detected tRFs of different lengths derived from both the tRNA^ArgU^ and tRNA^LysT^ corresponding to tRNA halves, tRF 5’ and tRF 3’. Interestingly, as observed in **Fig. 7C** the tRNA^LysT^ and tRNA^ArgU^, the 3’ fragment levels go down in the tRNA^ArgU^ and tRNA^LysT^ overexpressed strains respectively, when compared with the vector control. When we analysed further, we observed that upon overexpression of tRNA^LysT^ there was downregulation of tRNA^ArgU^ and interestingly, overexpression of tRNA^ArgU^ caused downregulation of tRNA^LysT^ (**Fig. S6**). This observation further supports the tRNA^LysT^ and tRNA^ArgU^ interaction *in vivo* and sequestration of tRNA^ArgU^ by tRNA^LysT^. Overall, these data provide yet another potential mechanism for tRNA interaction where mature tRNA-tRFs or tRF-tRFs may interact with each other (**Fig. 8**).

**Fig. 7:**
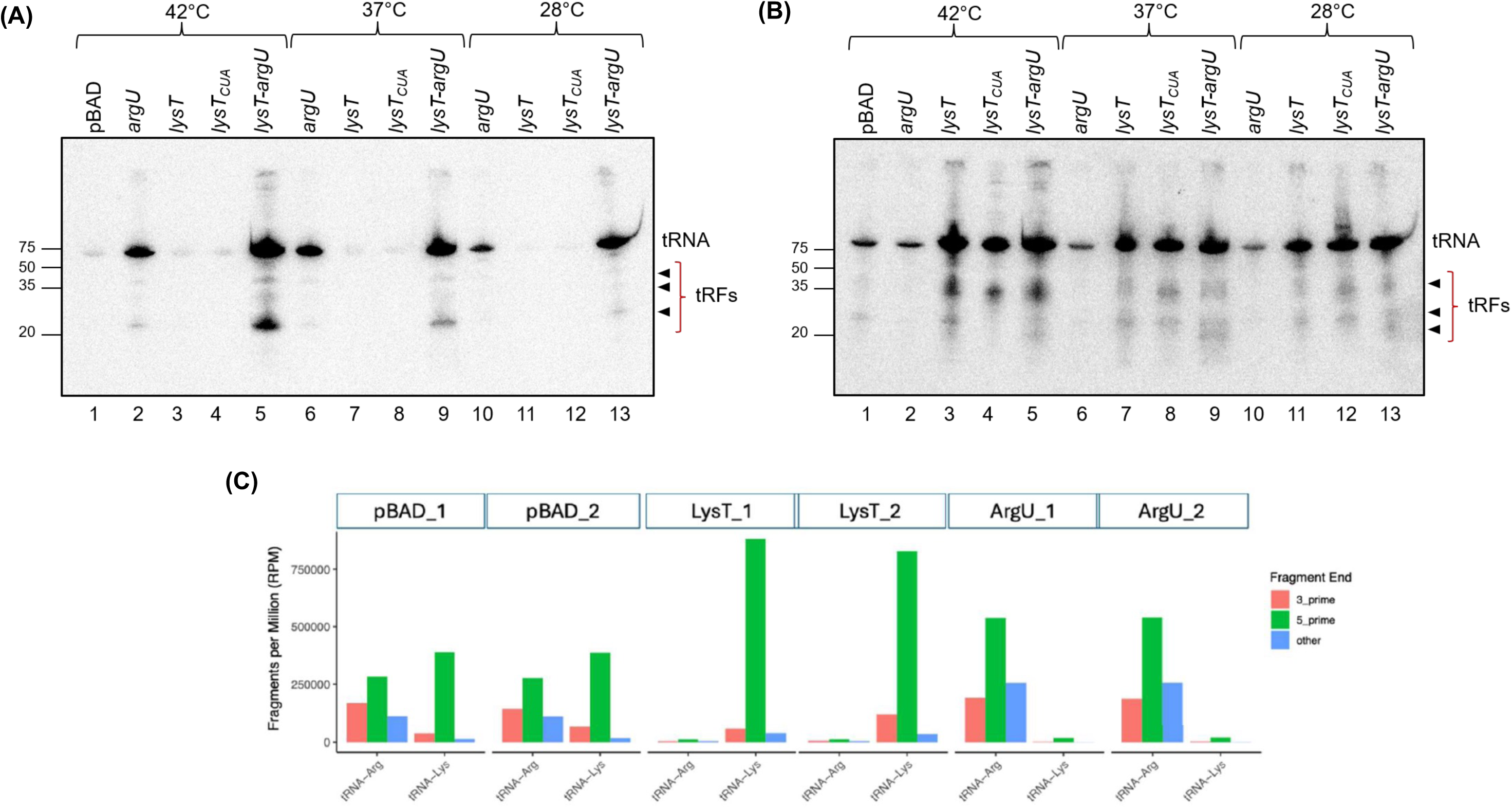
tRNA derived fragments (tRFs) detected by northern blotting. **(A)** tRNA^ArgU^ 5′ end and **(B)** tRNA^LysT^ and tRNA^ArgU^ 3′ end at different temperatures. tRFs obtained from tRNA overexpressed in BW25113 strain grown in LB media at 42, 37 and 28 °C. Arrowhead depicts tRFs. **(C)** Grouped bar plots show the distribution of normalized fragment counts per tRNA gene and fragment end type obtained from small RNA sequencing analysis.

**Fig. 8:**
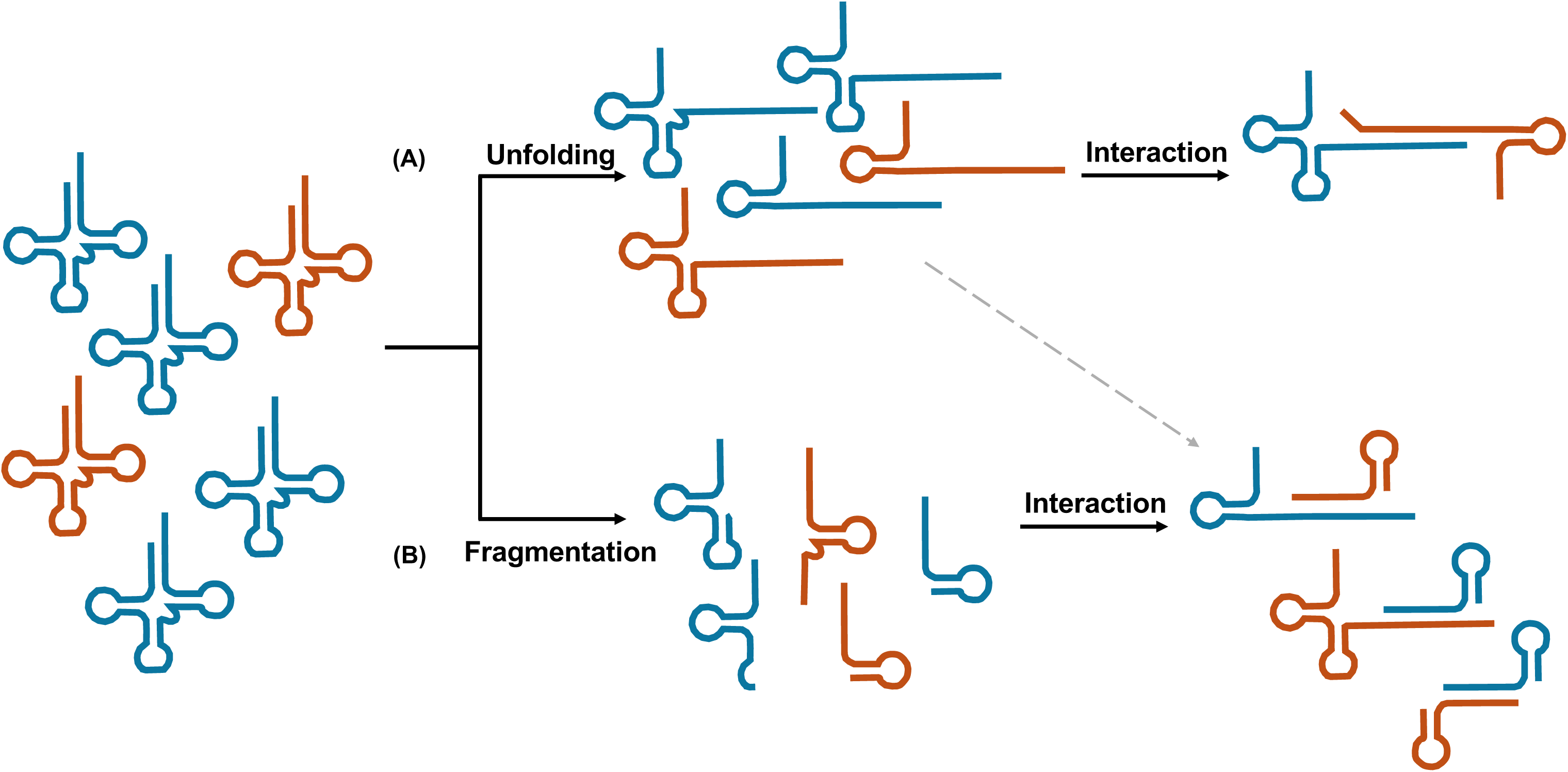
Schematic diagram showing the proposed model for tRNA^ArgU^ (in orange) and tRNA^LysT^ (in blue) interaction *in vivo*. (**A**) Mature tRNA may undergo denaturation or attain non-canonical conformation under some stress conditions and facilitate tRNA^ArgU^-tRNA^LysT^ heterodimer formation. Alternatively, the action of chaperone proteins, like in the case of HIV tRNA^Lys3^ may unfold the tRNA, and help tRNA-tRNA heterodimer formation. (**B**) tRNA may interact with each other through tRNA-derived fragments (tRFs). Both the tRNA^ArgU^ and tRNA^LysT^ form fragments which may interact with mature tRNAs or tRFs.

## Discussion

Studies focusing on the sequences and structures of tRNAs, and the role of complementary sequences within and between tRNAs, continue to provide important leads into their evolution and functional diversities. We analysed bacterial (*E. coli*) tRNA sequences and found tRNA pairs with significant complementarity between them. We chose the tRNA^LysT^ and tRNA^ArgU^ pair for our investigation because of two main reasons; firstly, the RNAup analysis revealed a significant potential interaction between tRNA^LysT^ and tRNA^ArgU^, which maps to TΨC arm, with a predicted binding free energy of −8.64 kcal/mol, signifying a thermodynamically stable tRNA–tRNA duplex (**Fig. 1; Table: S1, S2**). Secondly, while tRNA^LysT^ is an abundant tRNA that decodes the AAA and AAG codons, tRNA^ArgU^ is a rare tRNA that decodes the AGA codon and, less frequently, the AGG codon (Chen & Texada, 2006). Our *in vitro* analysis offers strong evidence that the tRNA^LysT^ and tRNA^ArgU^ interact with each other, supporting the possibility of intermolecular interactions between complementary tRNAs (**Figs. 2** and **3**). Furthermore, to determine whether this interaction has physiological significance, we designed experiments to assess the effects of these tRNA pairs *in vivo*. Growth studies showed that while the overexpression of tRNA^LysT^ causes slow growth at low temperatures, the overexpression of tRNA^ArgU^ showed little effect on growth. These observations correlate well with the levels of tRNA^LysT^ and tRNA^ArgU^ in the cell (**Fig. 4**), and are consistent with a previous report showing that the accumulation of some tRNAs is toxic to the cell (Zimmerman et al., 2018).

To investigate the *in vivo* effects of altered abundances of tRNA^LysT^ and tRNA^ArgU^, we used two independent assays: one based on the λ_imm_-P22 phage growth on the *ssrA* mutant strain and the other based on the toxicity of the AGA minigene. In the *ssrA*^AGA^ mutant strain background, tRNA^LysT^ overexpression caused toxicity and inhibition of the phage growth (**Fig. 5A**); and in AGA minigene background, tRNA^ArgU^ over-expression alleviates the growth inhibition to significant extent (**Fig. 6**). These results provide us with an evidence of tRNA^LysT^-tRNA^ArgU^ interaction and tRNA^ArgU^ sequestration by tRNA^LysT^.

Earlier reports (Liu et al., 2017; Loehr & Keller, 1968; Yang et al., 1972), have shown tRNA-tRNA interactions (hetero or homodimer) in bacterial cells. In this study, we focused on pair of tRNAs where tRNA^LysT^ is an abundant species and tRNA^ArgU^ is a rare species, and which show heterodimer formation *in vitro*. The observations from the minigene based *in vivo* experiments are consistent with the interaction of tRNA^LysT^ (abundant) and tRNA^ArgU^ (rare). However, the physiological cues of how these tRNAs interact with each other remains unclear. We propose two pathways for this interaction to happen. First, the mature tRNAs may unfold or attain non-canonical conformations under the stressful temperatures/ionic conditions facilitating their interaction to allow regulation of protein synthesis (**Fig. 8A**). Although the cloverleaf secondary structure is natural for the majority of tRNAs, databases exhibit sequence variations that do not align with the standard cloverleaf (Giegé et al., 2012). Fundamentally, the folding of secondary structures is not solely reliant on sequence but is also affected by biophysical and chemical factors. And the Mg^2+^-dependent alternative folding of hairpin and cloverleaf in *E. coli* tRNA^Glu^ was characterized (Madore et al., 1999). This suggests that any changes in the environmental condition around the tRNA may allow it to obtain a non-canonical configuration, which in turn makes it feasible to interact or form a dimer with another tRNA. We have also shown the possible tRNA^LysT^ and tRNA^LysT^ dimer conformation by using *bifold* server which predict the lowest hybrid free energy conformation (**Fig. S7 and S8**). Another possible way of how tRNA may unfold is by the action of chaperone proteins. It is reported that chaperone proteins can help in tRNA folding or unfolding (Keffer-Wilkes et al., 2016b, 2020; Vakiloroayaei et al., 2017); and help in maintaining RNA-RNA interactions (Katsuya-Gaviria et al., 2022; Melamed et al., 2020; Park et al., 2021). We assume that tRNA^LysT^ and tRNA^ArgU^ may unfold with the help of some chaperone proteins to interact/heteromerize with each other. This mechanism may be similar to what was earlier reported for the viral tRNA^Lys3^, which unfolds from canonical cloverleaf structure to act as a primer for the viral replication (Kleiman, 2002; Mak & Kleiman, 1997).

Another possibility of regulation could be presented through the interaction through the tRNA-derived fragments (tRFs). We observed tRFs from both tRNA^ArgU^ and tRNA^LysT^ (**Fig. 7**) raising the possibility that these fragments contribute to the observed regulatory effects. Small RNA sequencing analysis shows the presence of the tRNA^ArgU^ and tRNA^LysT^ fragments. It has been reported that tRFs from different tRNAs do form homodimer, heterodimer or even tetramers (Lyons et al., 2017; Tosar et al., 2018). Therefore, we propose that tRNA fragments might interact either with the full length tRNAs or with each other (**Fig. 8B**). We observed that 5’ fragments dominate in the tRNA overexpressed strains. Interestingly, while tRNA^LysT^ overproduction increases tRNA^LysT^ fragments it decreases the tRNA^ArgU^ fragments (**Fig. 7C**). Likewise, while the tRNA^ArgU^ overexproduction increases tRNA^ArgU^ fragments it decreases the tRNA^LysT^ fragments. These observations suggests that the 3’ fragments of tRNA^LysT^ and tRNA^ArgU^ could form dimers which in turn may lead to their degradation. Overall, these data suggest that such interactions could potentially modulate tRNA availability, stability, or function and represent an additional layer of post-transcriptional regulation.

Although *lysT* overexpression markedly increased the abundance of tRNA^Lys^, polysome profiling revealed a reduction in polysome formation, indicating attenuation of global translation (**Fig. S9**). In contrast, *argU* overexpression increased tRNA^ArgU^ abundance without altering polysome profiles, indicating that elevated tRNA^ArgU^ levels are better tolerated and do not significantly perturb global translation. These data (along with the alterations in the tRF^ArgU^ levels of upon overexpression of tRNA^Lys^ and vice versa) also support the idea that tRNA^LysT^ interacts with tRNA^ArgU^ and regulates translation. However, a limitation of our study is that we were unable to show *in vivo*, a direct physical interaction between tRNA^LysT^ and tRNA^ArgU^ despite attempts to investigate for such an interaction. More sensitive methods may be helpful in detecting tRNA^LysT^ and tRNA^ArgU^ interactions *in vivo*. Nonetheless, multiple lines of evidence we provide here support an *in vivo* interaction between tRNA^LysT^ and tRNA^ArgU^, especially when tRNA^LysT^ is overproduced in the cell. It would be interesting to check more such tRNA pairs for their interactions and their role in the regulation of cellular functions.

## Materials and Methods

### Media and chemicals used

Bacteria were grown in Luria-Bertani (LB) broth, or LB agar plates containing 1.8% bacto agar (Difco) supplemented with ampicillin (Amp; 100 µg/mL), kanamycin (Kan; 25 µg /mL), tetracycline (Tet; 7.5 µg/mL), arabinose or glucose as required (in **Fig. S1**). Chemicals, reagents, and enzymes were obtained from Thermo Scientific, Sigma (USA), New England Biolabs (USA) and SRL (India). Radioisotopes were acquired from American Radiolabelled Chemical (USA) and Board of Radiation and Isotope Technology (India).

### Strains, plasmids, DNA oligomers and growth analysis

The bacterial strains and plasmids used in this work are listed in Tables S3 and S4, respectively. DNA oligomers listed in Table S5 were synthesised by Sigma (India).

### Comparison of tRNA gene sequences

All *E. coli* tRNA sequences were obtained from the tRNAScan genomic tRNA database (http://gtrnadb.ucsc.edu/) (Chan & Lowe, 2009, 2016). Pairwise comparisons between the sequences were performed using the RNAup server (Gruber et al., 2008). The free energy of tRNA-tRNA interactions was calculated using both RNAup server and *bifold* algorithm implemented in the RNAstructure web server (Reuter & Mathews, 2010).

### tRNA synthesis by *in vitro* transcription

For *in vitro* transcription of tRNAs, we followed an earlier protocol (Korencić et al., 2002). Briefly, 99 bp long tRNA transcription template with a T7 promoter upstream of the *argU* tRNA sequence was obtained by annealing two oligomers, ArgU_T7Fp and ArgU_T7Fp (for sequences see Table S1) with overlapping sequences (19 bp overlap). Similarly, for tRNA^LysT^ and tRNA^ValU^, DNA oligomers LysT_T7Fp & LysT_T7Fp and ValU_T7Fp & ValU_T7Fp, respectively were used. Reaction (50 µL) with about 500 pmol of each oligomer in 1X Klenow polymerase buffer was heated to 95 °C for 5 min and allowed to gradually cool to 25 °C. Then, 250 μM of dNTPs and 10 U of Klenow DNA polymerase were added to the mixture and extension was allowed to proceed for 30 min at 37 °C. The reaction products were extracted, ethanol precipitated and resuspended in RNase free water. *In vitro* transcription was carried out using Thermos Scientific 1X transcription buffer, 2 mM NTPs (either radiolabelled ꭤ-GTP or unlabelled GTP was used), 50 U Ribolock RNase inhibiter, 100 U T7 RNA polymerase and 2 µg of template were incubated for 2 h at 37 °C. The template DNA was removed from the reaction by 4 U DNase I digestion. The reaction product was phenol extracted, ethanol precipitated and resuspended in RNase free sterile water. The products were analysed on 8 M, 15% urea PAGE. Use of radiolabelled ꭤ-GTP provided radiolabelled tRNA transcripts.

### *In vitro* binding of tRNAs

*In vitro* tRNA binding/annealing assay was set up in a reaction containing 2 µg of either tRNA^LysT^- tRNA^ArgU^ or tRNA^LysT^-tRNA^ValU^ or tRNA^ArgU^-tRNA^ValU^ (where either of them was radiolabelled) in 0.5X SSC buffer, was incubated at 95 °C for 5 min and allowed to cool down to 25 °C. The reaction mixture was UV crosslinked and analysed on 15% native PAGE at 4 °C followed by northern blotting. A BioImage Analyzer (FLA5100, Fujifilm), was used to image the gel exposed on phosphor-imaging screen. The binding assays were also performed in the presence of different concentrations of the competitor DNA oligomers (Table S3).

### Generation of pBAD*lysT* and pBAD*argU* constructs

LysT and ArgU tRNA genes were amplified by PCR from *E. coli* genomic DNA using LysT_Fp_NcoI & LysT_Rp_EcoRI and ArgU_Fp_EcoRI & ArgU_Rp_HindIII primers, respectively. LysT and ArgU tRNA amplicons were digested with NcoI & EcoRI and EcoRI & HindIII restriction enzymes respectively. Similarly, pBAD vector was also digested with same sets of restriction enzymes. The tRNA amplicons were cloned in this vector to result in pBAD*lysT* and pBAD*argU*. LysT and ArgU tRNA genes were also cloned in the pTrc*AGA* minigene constructs at BstZ17I site by blunt-end cloning, and confirmed by DNA sequencing.

### Generation of pACDH*ssrA*^WT^ and pACDH*ssrA*^AGA^ constructs

Wild type *ssrA* gene was cloned into SacI and NcoI restriction site of pACDH vector to generate pACDH*ssrA*^WT^ construct. The resume codon of *ssrA* gene was mutated from GCA to AGA by inverse PCR on pACDH*ssrA*^WT^, using ssrA_AGA_SDM_Fp and ssrA_AGA_SDM_Rp primers to generate pACDH*ssrA*^AGA^.

### Generation of BW25113Δ*ssrA::kan* strain

BW25113Δ*ssrA*::*kan* strain was generated by P1 phage mediated transduction. BW25113 strain was transduced with P1 lysate raised on TG1Δ*ssrA::kan* strain to generate BW25113Δ*ssrA::kan* strain

### Growth curve analysis

Four biological replicates of each strain were inoculated in LB with desired antibiotic(s) and grown at 37 ^°^C overnight. For growth analysis, the overnight cultures were diluted thousand-fold (10^-3^ dilution) in LB media, and 200 μL of it was seeded in honeycomb plates for automated growth in BioScreen C at desired temperature, and OD_600_ was measured every hour. OD_600_ values (with standard mean) were plotted against time using GraphPad Prism v10.2.1.

### Total tRNA isolation, native PAGE and northern blot

Total tRNA isolation, native PAGE analysis and northern blotting were performed as described (Shetty et al., 2016) using probes as mentioned in Table S5.

### Phage plating assay

The assays were performed as before (Kapoor et al., 2011; Singh et al., 2008). Briefly, bacterial cultures were grown till late log phase in LB with 10 mM MgSO_4_, 1% maltose, and induced with 0.2 mM IPTG. A culture pellet of 1 OD_600_ resuspended in 100 µl media was taken, mixed with 10^-6^ dilution of hybrid λ_imm_-P22 phage lysate, incubated for 15 min at 37 °C, mixed with top agar (0.8%), poured onto LB agar plates with the desired antibiotics, incubated at 28 °C for 8-12 h and scored for the plaque numbers, and their sizes using Image Lab v6.1 software.

### tRNA-derived fragment detected by northern blotting

Total tRNA was separated on 8M, 15% urea PAGE followed by northern blotting using specific radiolabelled probes (Table S3) designed against 3′ end of tRNA^LysT^ and tRNA^ArgU^.

### RNA library preparation and sequencing

RNA library preparation and small RNA sequencing was done from Chromosome Labs Pvt. Ltd. Briefly, 5 µg of RNA was processed using rtStar tRF and tiRNA Pretreatment kit (Cat. no. AS-FS005, Arraystar Inc.), according to the manufacturer’s protocol with minor modifications in RNA cleanup. Libraries were prepared for small RNA by using SMARTer smRNA-Seq Kit for Illumina (Cat. No. 635031, Takara Bio), according to the manufacturer’s protocol. The libraries were sequenced on NextSeq2000 (Illumina Inc.) using PE50 sequencing chemistry.

### RNA sequencing data analysis

The raw data obtained from Illumina sequencing were recorded in FASTQ files. Quality of the raw reads was assessed using FastQC (Andrews, S. 2010), and cutadapt (Martin, 2011) was used to trim and filter the raw reads. The adapters were removed, and the reads were dropped if the average quality of the bases dropped below 30. As the target was tRNA and its fragments, the minimum read length cutoff was set as 15 bp. The filtered reads were mapped to *E. coli* BW25113 strain K-12 (NZ_CP009273.1) genome using bowtie2 (Langdon, 2015) with parameters adapted for very short reads mapping (--local --very-sensitive-local -N 1 -L 15 - iS,1,0.75 --mp 2,2). The aligned and sorted bam files were generated using samtools and reads were counted against the reference annotation file using feature Counts (Li et al., 2009). Reads mapped to all tRNAs were extracted from the bam files and then their quantification was carried out using bedtools and analysed using R. The comparison of read length distribution was carried out between targeted tRNA (tRNA^LysT^ and tRNA^ArgU^) with other tRNAs. A histogram comparing the two tRNAs was plotted, and the mapping was visualized using IGV (Liao et al., 2014).

## ACKNOWLEDGEMENTS

The authors acknowledge laboratory colleagues for their critical comments on the manuscript, and Dr. Neeraja Krishnan for her initial observations on tRNA complementarities.

